# Group B rotavirus encodes a functional fusion-associated small transmembrane (FAST) protein

**DOI:** 10.1101/639070

**Authors:** Julia R. Diller, Helen M. Parrington, John T. Patton, Kristen M. Ogden

**Author notes:** Address correspondence to Kristen M. Ogden. John Patton, Department of Biology, Indiana University, Bloomington, Indiana, USA.

## Abstract

Rotavirus is an important cause of diarrheal disease in young mammals. Group A rotavirus (RVA) causes most human rotavirus diarrheal disease and primarily affects infants and young children. Group B rotavirus (RVB) has been associated with sporadic outbreaks of human adult diarrheal disease. RVA and RVB are predicted to encode mostly homologous proteins but differ significantly in the proteins encoded by the NSP1 gene. In the case of RVB, the NSP1 gene encodes two putative protein products of unknown function, NSP1-1 and NSP1-2. We demonstrate that human RVB NSP1-1 mediates syncytia formation in cultured human cells. Based on sequence alignment, NSP1-1 from groups B, G, and I contain features consistent with fusion-associated small transmembrane (FAST) proteins, which have previously been identified in other *Reoviridae* viruses. Like some other FAST proteins, RVB NSP1-1 is predicted to have an N-terminal myristoyl modification. Addition of an N-terminal FLAG peptide disrupts NSP1-1-mediated fusion, consistent with a role for this fatty-acid modification in NSP1-1 function. NSP1-1 from a human RVB mediates fusion of human cells but not hamster cells and, thus, may serve as a species tropism determinant. NSP1-1 also can enhance RVA replication in human cells, both in single-cycle infection studies and during a multi-cycle time course in the presence of fetal bovine serum, which inhibits rotavirus spread. These findings suggest potential yet untested roles for NSP1-1 in RVB species tropism, immune evasion, and pathogenesis.

**IMPORTANCE:** While group A rotavirus is commonly associated with diarrheal disease in young children, group B rotavirus has caused sporadic outbreaks of adult diarrheal disease. A major genetic difference between group A and B rotaviruses is the NSP1 gene, which encodes two proteins for group B rotavirus. We demonstrate that the smaller of these proteins, NSP1-1, can mediate fusion of cultured human cells. Comparison with viral proteins of similar function provides insight into NSP1-1 domain organization and fusion mechanism. Our findings are consistent with an important role for a fatty acid modification at the amino terminus of the protein in mediating its function. NSP1-1 from a human virus mediates fusion of human cells, but not hamster cells, and enhances rotavirus replication in culture. These findings suggest potential, but currently untested, roles for NSP1-1 in RVB species tropism, immune evasion, and pathogenesis.

## INTRODUCTION

Rotaviruses are members of the *Reoviridae* family of nonenveloped viruses with segmented dsRNA genomes and causative agents of diarrheal disease in many young mammals, including humans. Adults are often resistant to rotavirus diarrheal disease. Acquired immunity, particularly local and systemic antibodies, plays an important role in protection from rotavirus disease, and immunity appears to increase with repeated infection or immunization (1). However, innate immunity also may contribute to rotavirus disease, and rotaviruses have been shown to antagonize innate signaling pathways using multiple distinct mechanisms (1–3).

Rotaviruses are currently classified into eight species, A-I, which further resolve into two major clades (https://talk.ictvonline.org/) (4–6). One clade contains species A, C, D, and F, and the other contains species B, G, H, and I. The majority of human rotavirus diarrheal disease occurs in infants and young children and is associated with rotavirus species A (RVA) (1). RVA also causes diarrheal disease in birds and numerous mammals, though subsets of RVA genotypes are associated with specific hosts (7). RVB, RVC, RVH, and RVI have been detected mostly in domesticated mammals, while RVD, RVF, and RVG have been detected in birds. However, instances of zoonotic transmission of rotaviruses and their gene segments, particularly between humans and domesticated animals, have been documented (8, 9). Although some factors, such as lack of appropriate attachment and entry machinery or adaptive immune cross-protection, are known to impose barriers, factors permitting or limiting rotavirus zoonotic transmission remain incompletely understood.

Evidence of RVB infection has been commonly detected in diarrheic pigs (10–12), and RVB has been associated with sporadic outbreaks of diarrheal disease in humans (13–16). The first reported human RVB outbreak occurred in China from 1982-1983, ultimately affecting more than a million people with cholera-like diarrhea (17–21). While RVB disease symptoms resemble those of RVA gastroenteritis, RVB causes disease primarily in adults rather than pediatric populations (22). Studies suggest there is low-level RVB seroprevalence in humans (23–25). RVB outbreaks in humans are not thought to be caused by viruses directly transmitted from animals; rather, phylogenetic analysis of RVB sequences suggests viruses affecting humans and other animals are distinct (26). Factors contributing to the capacity of these viruses to cause disease in adults remain unknown.

Like RVA, RVB have genomes containing 11 segments of dsRNA. Based on sequence alignment and structure prediction, 10 of the 11 RVB segments encode proteins with RVA homologs (27, 28). However, the segment encoding RVA innate immune antagonist NSP1 differs significantly. For most rotaviruses in the clade containing RVB, including RVG and RVI, the NSP1 gene segment contains two overlapping ORFs whose encoded protein products have little predicted sequence or structural homology with known proteins (29). The first RVB ORF, NSP1-1, is predicted to encode a product approximately 100 amino acids in length (Fig. 1A), though the protein product has not been shown to be expressed. The length and predicted structural features of NSP1-1 are reminiscent of fusion-associated small transmembrane (FAST) proteins.

**Figure 1.**
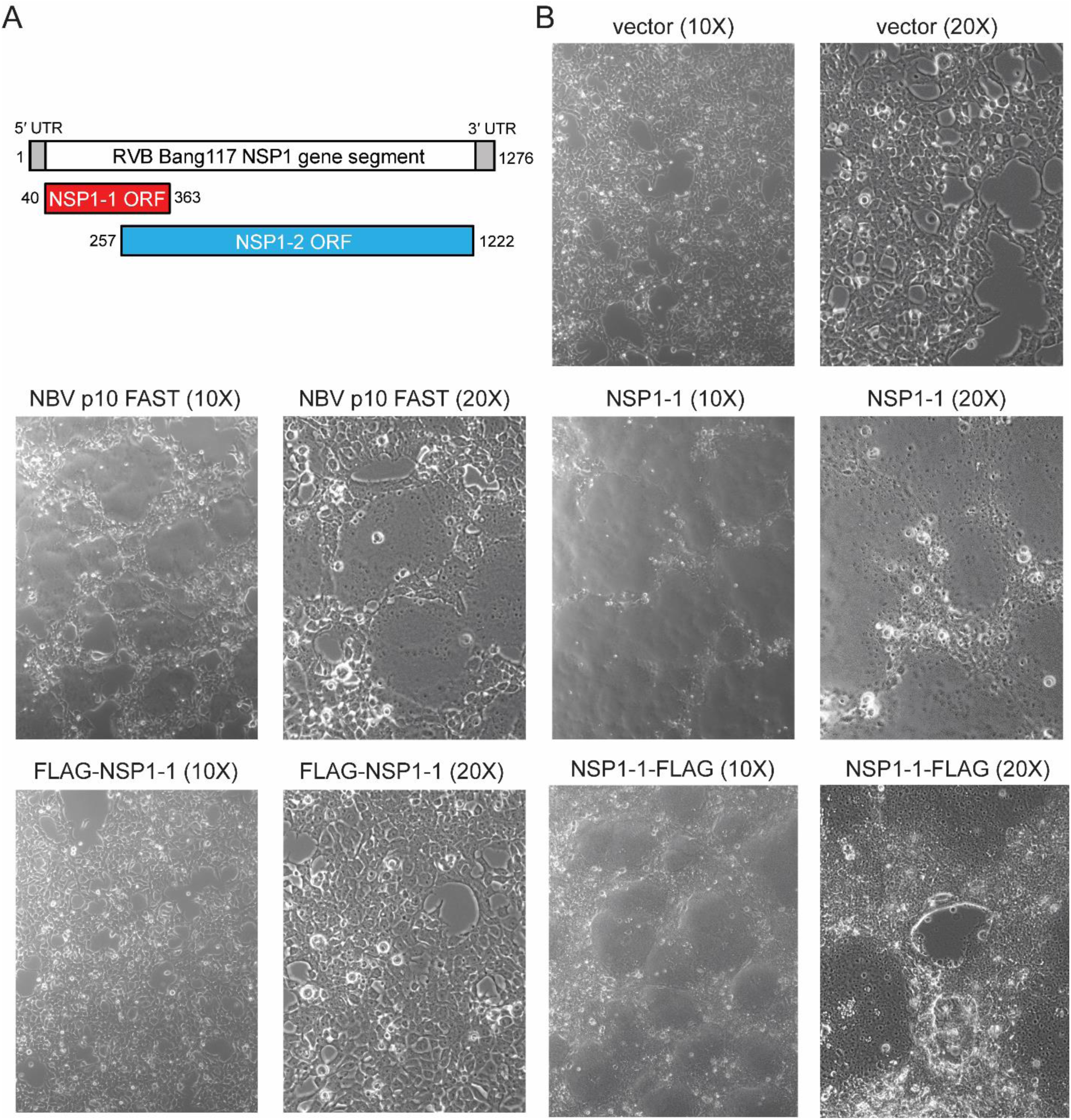
RVB Bang117 NSP1-1 mediates syncytia formation in 293T cells. (A) Schematic showing the organization of RVB Bang117 NSP1 gene segment, including untranslated regions (UTRs) and two putative ORFs. (B) Differential interference contrast images of 293T cells transfected with vector alone or plasmids encoding NBV p10 FAST or RVB Bang117 NSP1-1 in its untagged form or with an N- or C-terminal FLAG tag. Representative images are shown. Plasmids used for transfection and objective lens magnification are indicated.

FAST proteins are a family of small, bitopic plasma membrane proteins that mediate cell-cell fusion and syncytium formation (reviewed in (30, 31)). These nonstructural viral proteins are encoded by fusogenic members of the *Aquareovirus* and *Orthoreovirus* genera of the nonenveloped *Reoviridae* virus family. There are multiple types of FAST proteins, and they range in size from approximately 90-200 amino acids. Each contains three modular domains: a small, N-terminal extracellular domain that is often acylated, a transmembrane domain, and a C-terminal cytoplasmic tail containing a polybasic motif. Additional putative functional motifs have been identified and vary among FAST protein family members. FAST proteins are nonstructural proteins that are expressed following virus entry, viral mRNA transcription, and translation. Membrane fusion occurs in closely apposed cells. FAST protein clustering and interactions with the opposing lipid bilayer, including insertion of fatty acid moieties or hydrophobic residues, favors lipid mixing and membrane curvature, leading to pore formation. Following pore formation, cellular proteins, including annexin 1 and actin promote pore expansion and, thereby, syncytia formation.

In the current publication, we provide evidence that RVB NSP1-1 is a FAST protein that is capable of mediating syncytia formation in some, but not all, mammalian cell types. Based on sequence alignment, we suggest that other rotavirus species, RVG and RVI, also may encode functional FAST proteins. We demonstrate that the N terminus is required for NSP1-1-mediated fusion and provide experimental support for a role of NSP1-1 in viral replication and spread. These findings have potential implications for the role of NSP1-1 in host immune evasion and human RVB disease.

## RESULTS

### The N terminus is required for RVB NSP1-1-mediated fusion in 293T cells

To test the hypothesis that NSP1-1 is a FAST protein, we transfected human embryonic kidney 293T cells with a pCAGGS vector, pCAGGS encoding the fusogenic Nelson Bay orthoreovirus (NBV) p10 FAST protein (32), or pCAGGS encoding codon-optimized human RVB Bang117 NSP1-1, permitted expression for 24 h, then examined cell morphology using differential interference contrast microscopy. Transfection with NBV p10, a known FAST protein (32–35), or with RVB NSP1-1 changed the cell morphology from individually distinct cells to a monolayer pockmarked by smooth oval-shaped regions lacking defined cell edges, which likely represent syncytia (Fig. 1B). These observations suggest that, like NBV p10, RVB NSP1-1 can mediate cell-cell fusion.

To enable detection of RVB NSP1-1, we engineered a FLAG peptide at the N or C terminus. Following transfection of 293T cells with pCAGGS encoding tagged forms of RVB NSP1-1, we found that C-terminally tagged NSP1-1 (NSP1-1-FLAG) mediated morphological changes resembling syncytia in the cell monolayer (Fig. 1B). Cells transfected with plasmids encoding N-terminally tagged NSP1-1 (FLAG-NSP1-1), however, were morphologically indistinguishable from vector-transfected cells. This finding suggests that the N terminus plays an important role in RVB NSP1-1-mediated cell morphological changes.

To gain insight into RVB NSP1-1 localization and cell morphological changes, we transfected 293T cells with pCAGGS encoding FLAG tagged NSP1-1, waited 24 h to permit protein expression, fixed and stained the cells to detect FLAG and nuclei, and imaged them using confocal microscopy. FLAG-NSP1-1 was typically expressed in the cytoplasm of individual, or sometimes adjacent, cells (Fig. 2A). In Z-stacks, FLAG-NSP1-1 was detected in the cytoplasm at the level of the nucleus, and individual stained cells were distinct (Fig. 2C). In striking contrast to FLAG-NSP1-1, NSP1-1-FLAG was detected in clusters containing many nuclei (Fig. 2B). When looking through a Z-stack, NSP1-1-FLAG was detected at and above the level of the nucleus, consistent with cellular and plasma membrane localization, and the edges of individual stained cells were indistinguishable (Fig. 2D).

**Figure 2.**
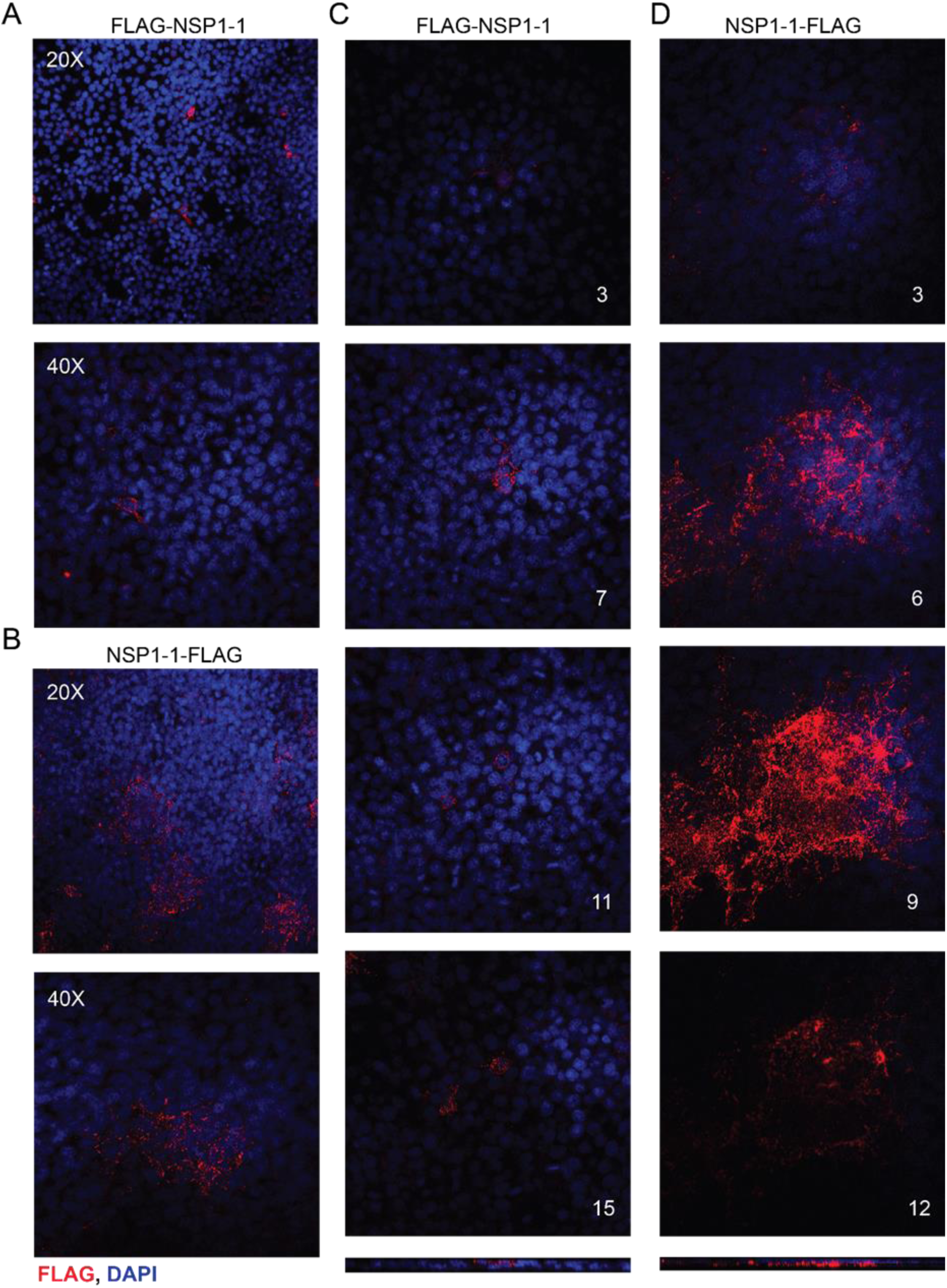
RVB Bang117 NSP1-1 localization in 293T cells. 293T cells were transfected with plasmids encoding RVB Bang117 NSP1-1 with an N- or C-terminal FLAG tag, as indicated. Cells were fixed and stained with antibodies to detect FLAG (red) or nuclei (blue) and imaged using confocal microscopy. (A-B) Images from a single confocal plane of 293T cells transfected with plasmids encoding FLAG-NSP1-1 (A) or NSP1-1-FLAG (B) taken using the 20X or 40X objective, as indicated. (C-D) Confocal images from comparable focal planes in a Z-stack taken using the 40X objective for 293T cells transfected with plasmids encoding FLAG-NSP1-1 (B) or NSP1-1-FLAG (C). Z-section number is indicated; numbers increase coincident with distance from the adherent surface of the monolayer. Bars at the bottom represent orthogonal views through the Z-stack, approximately at the center of the images.

### Shared features of RVB NSP1-1 and *Reoviridae* FAST proteins

The similarity in protein size and cell morphological changes induced upon RVB NSP1-1 expression suggested that it may be a FAST protein. To gain insight into the relationships between rotavirus NSP1-1 proteins and *Reoviridae* FAST proteins, we constructed a maximum likelihood (ML) tree using the sequences of representative RVB, RVG, and RVI NSP1-1 proteins and orthoreovirus and aquareovirus FAST proteins (Fig. 3A). We found that aquareovirus, and ARV/NBV FAST proteins each formed a clade supported by strong bootstrap values that clustered distinctly from the rotavirus NSP1-1 proteins and BRV p15, BroV p13, and RRV p14 FAST proteins. While they did not cluster together strongly, RVB and RVG NSP1-1 proteins clustered most closely with BRV p15, whereas RVI NSP1-1 proteins clustered more closely with RRV p14.

**Figure 3.**
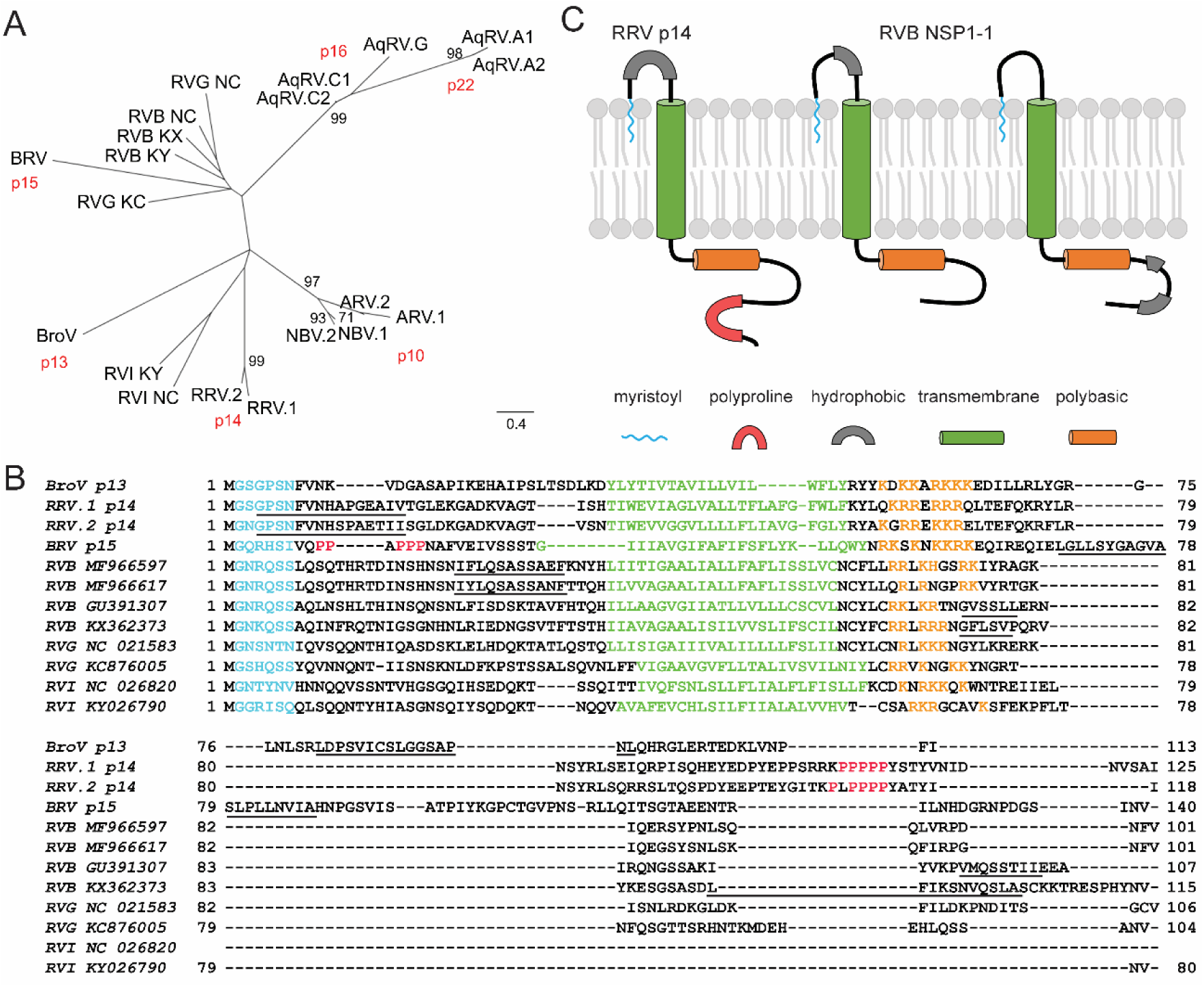
Conserved features of *Reoviridae* FAST proteins. (A) Maximum likelihood tree showing relationships among *Reoviridae* FAST and rotavirus NSP1-1 proteins. Abbreviations are: AqRV.A1, Atlantic salmon aquareovirus; AqRV.A2, Turbot aquareovirus; AqRV.G, American grass carp aquareovirus; AqRV.C1,Golden shiner aquareovirus; AqRV.C2, Grass carp aquareovirus; ARV.1, Avian orthoreovirus 176; ARV.2, Psittacine orthoreovirus; NBV.1, Nelson Bay orthoreovirus; NBV.2, Melaka orthoreovirus; BroV, Broome orthoreovirus; RRV.1, Python orthoreovirus; RRV.2, Green bush viper orthoreovirus; BRV, Baboon orthoreovirus; RVB KY, Rotavirus B strain RVB/Goat-wt/USA/Minnesota-1/2016; RVB KX, Rotavirus B strain RVB/Pig-wt/VNM/12089_7; RVB NC, Human rotavirus B strain Bang373; RVG KC, Rotavirus G pigeon/HK18; RVG NC, Rotavirus G chicken/03V0567/DEU/2003; RVI NC, Rotavirus I strain KE135/2012; RVI KY, Rotavirus I cat. Scale, in amino acid substitutions per site, is indicated. (B) Alignment of selected *Reoviridae* FAST and rotavirus NSP1-1 proteins. Abbreviations for FAST proteins are as in (A). Rotavirus species (RVB, RVG, or RVI) and accession number are indicated. Predicted N-myristoylation motifs are colored cyan, transmembrane helices are colored green, polybasic regions are colored orange, polyproline regions are colored red, and hydrophobic regions are underlined. (C) Cartoon models highlighting the predicted features and membrane topology for the RRV p14 FAST protein and for RVB NSP1-1. Features and models of BroV p13, RRV p14, and BRV p15 shown in (B) and (C) are based on previously published work (31).

To gain insight into sequence and structural features of rotavirus NSP1-1 proteins, we used software to scan for sequence motifs and constructed amino acid alignments with the most closely clustering FAST proteins from the ML tree (Fig. 3A). Based on the PROSITE definition (PDOC00008), an N-myristoylation site was predicted at amino acids 2-7 in RVB NSP1-1 (Fig. 3B). Although there is a high false-positive prediction rate for N-myristoylation motifs, prediction at this precise location for every complete RVB, RVG, and RVI NSP1-1 sequence deposited in GenBank (as of 12/4/2018; Fig. S1) provides confidence in its legitimacy. BroV, RRV, and BRV FAST proteins are also known or predicted to be N-myristoylated (36–39). Using the TMHMM Server, we identified predicted transmembrane helices in RVB, RVG, and RVI sequences (Fig. 3B). In each case, the N terminus was predicted to be extracellular, while the C terminus was predicted to be cytoplasmic. For RVB Bang117 NSP1-1, the TM region was predicted to span amino acids 39-61. The N termini of the NSP1-1 proteins in the alignment were typically predicted to be shorter than those of the FAST proteins. Like the BroV, RRV, and BRV FAST proteins, each of the NSP1-1 proteins contained multiple basic residues shortly after the TM domain (Fig. 3B). However, fewer basic residues were present in NSP1-1 (4–5) than FAST (6–7) protein polybasic regions. Some RVB sequences contain short stretches of hydrophobic residues in the N-terminal domain, while others contain two short hydrophobic regions in the C-terminal domain (Fig. 3B). Analyzed RVG and RVI NSP1-1 proteins lacked strong hydrophobic signatures outside of the predicted transmembrane domain. NSP1-1 proteins were typically shorter than FAST proteins, by up to 36 amino acids, with most of the difference in length residing C terminal to the polybasic region (Fig. 3B). The motifs identified by sequence alignment and analysis (Fig. 3B) suggest models of RVB NSP1-1 in which the extracellular, myristoylated N-terminal domain precedes a single transmembrane domain and a short cytoplasmic tail containing a polybasic region, with some RVB NSP1 proteins containing a single hydrophobic region N-terminal to the transmembrane domain and others containing two hydrophobic regions C-terminal to the polybasic region (Fig. 3C).

### RVB NSP1-1 mediates syncytia formation in Caco-2 cells

Rotaviruses infect enterocytes in the human intestine. To determine whether RVB Bang117 NSP1-1 could mediate syncytia formation in a more biologically relevant cell type, we transfected human epithelial colorectal adenocarcinoma Caco-2 cells with pCAGGS encoding FLAG-tagged RVB NSP1-1. These cells can form polarized monolayers and morphologically and functionally resemble the enterocytes lining the small intestine. To achieve reasonable transfection efficiency, we transfected subconfluent Caco-2 monolayers, waited 24 or 48 h to permit protein expression, fixed and stained the cells to detect FLAG and nuclei, then imaged them using indirect immunofluorescence microscopy. Similar to observations made in 293T cells (Fig. 2A), we found that FLAG-NSP1-1 was primarily expressed in the cytoplasm of individual Caco-2 cells or small numbers of adjacent cells, whereas NSP1-1-FLAG was expressed mostly in the cytoplasm of clusters of cells containing multiple nuclei and lacking distinct cell boundaries (Fig. 4A). We quantified the numbers of single cells and clusters (at least three immediately adjacent FLAG-positive cells) present in wells of transfected Caco-2 cells at 24 h post transfection. Consistent with a role for the N terminus in cell-cell fusion, we found that there were significantly more FLAG-positive single cells in FLAG-NSP1-1-transfected than NSP1-1-FLAG-transfected wells (∼9 fold) and significantly more clusters present in NSP1-1-FLAG-transfected cells than FLAG-NSP1-1-transfected wells (∼3.5 fold) (Fig. 4B). Often, groups of FLAG-NSP1-1-transfected cells we identified as “clusters” appeared to be groups of three or four adjacent singly transfected cells. To quantify differences in cluster size between FLAG-NSP1-1-transfected and NSP1-1-transfected Caco-2, we measured cluster diameters and found that diameters of NSP1-1-FLAG clusters were significantly larger than those of FLAG-NSP1-1 (Fig. 4C). These findings suggest that C-terminally tagged RVB Bang117 NSP1-1 can mediate fusion of a cell type similar to that targeted during natural RVB infection.

**Figure 4.**
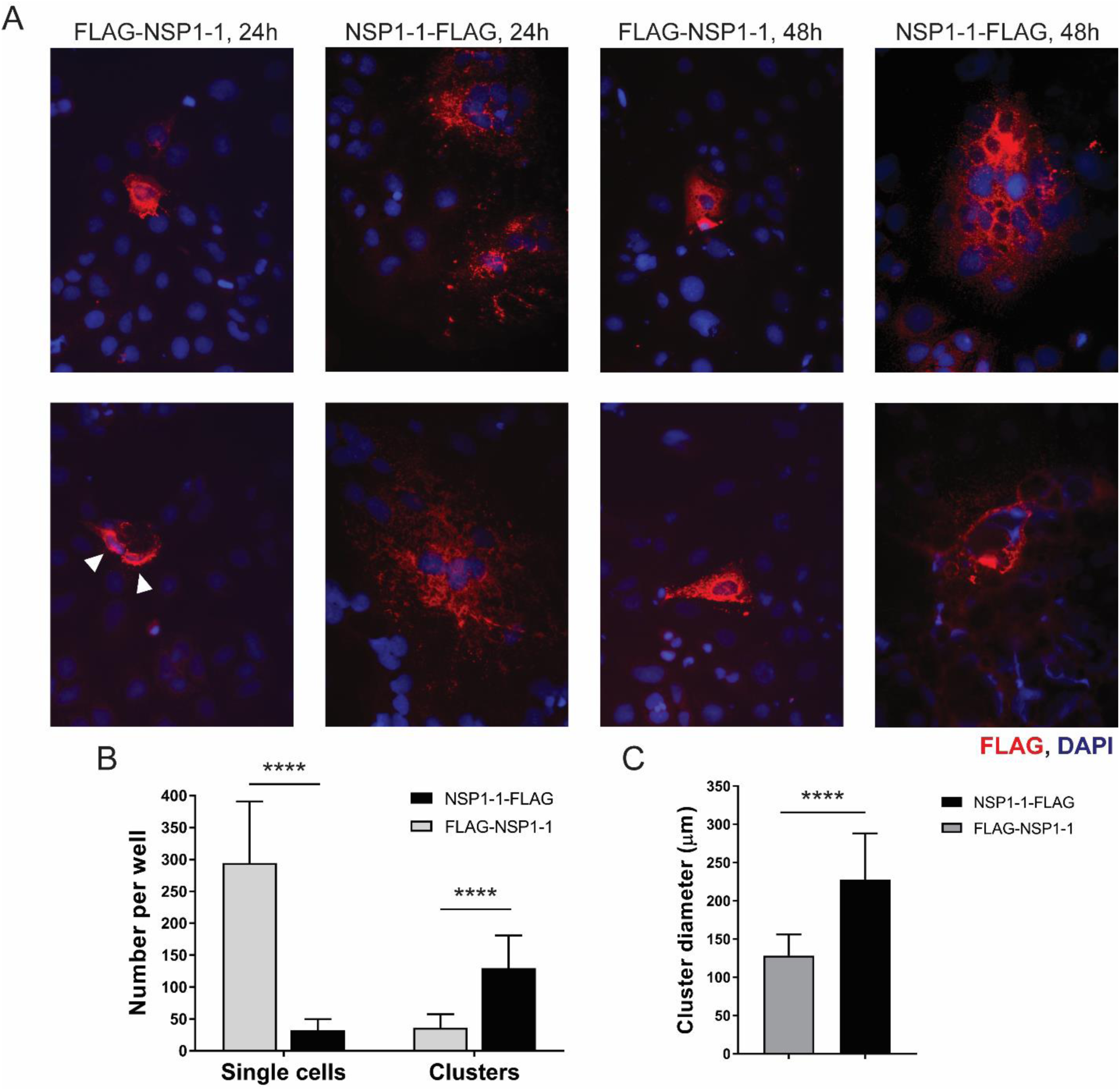
RVB Bang117 FAST mediates syncytia formation in Caco-2 cells. Caco-2 cells in 24-well plates were transfected with plasmids encoding RVB Bang117 NSP1-1 with an N- or C-terminal FLAG tag. Cells were fixed and stained with antibodies to detect FLAG (red) or nuclei (blue) and imaged using immunofluorescence microscopy at 24 and 48 h post transfection. (A) Representative images are shown. White triangles indicate adjacent FLAG-positive FLAG-NSP1-1-transfected cells. (B) The numbers of FLAG-positive clusters (3 or more adjacent cells) and FLAG-positive single cells per well were quantified in three wells per experiment in four independent experiments. *n* = 12. (C) The diameters of 20 FLAG-positive clusters per experiment in four independent experiments were quantified. *n* = 80. ****, *p* <0.0001 by unpaired *t* test.

### RVB NSP1-1 fails to mediate fusion in BHK cells

We next wanted to test RVB NSP1-1 function in the context of viral infection. NBV p10 enhances reovirus and rotavirus replication in baby hamster kidney cells expressing T7 RNA polymerase (BHK-T7 cells) (32). The reverse genetics system for simian rotavirus strain SA11 involves transfection of BHK-T7 cells with plasmids encoding the 11 rotavirus RNAs under the control of the T7 promoter, along with pCAGGS plasmids encoding viral capping enzymes, to enhance viral protein translation, and NBV p10, to enhance rotavirus replication and spread. We hypothesized that, if RVB NSP1-1 could mediate syncytia formation in BHK-T7 cells, it could functionally replace NBV p10 in rotavirus reverse genetics experiments. As a first step towards this goal, we transfected BHK-T7 cells with pCAGGS alone or pCAGGS expressing NBV p10, FLAG-NSP1-1, or NSP1-1-FLAG, waited 24 h to permit protein expression, fixed and stained the cells to detect FLAG and nuclei, and imaged them using differential interference contrast and indirect immunofluorescence microscopy. Transfection with pCAGGS encoding NBV p10 resulted in the appearance of balls or ring-like clusters of nuclei surrounded by regions of smooth cytoplasm nearly devoid of nuclei (Fig. 5B). Although many transfected cells detectably expressed NSP1-1-FLAG or FLAG-NSP1-1, no morphological differences were detected in comparison to vector-transfected cells, and all cells had distinct borders (Fig. 5A,C-D). A similar lack of morphological change was observed following transfection of BHK-T7 cells with untagged RVB NSP1-1 (not shown). Although localization was not examined in depth, NSP1-1-FLAG staining did not often diffusely fill the cytoplasm but appeared as perinuclear puncta, suggesting potential mislocalization (Fig. 5D). These findings suggest that RVB NSP1-1 is incapable of mediating cell-cell fusion in all mammalian cell lines.

**Figure 5.**
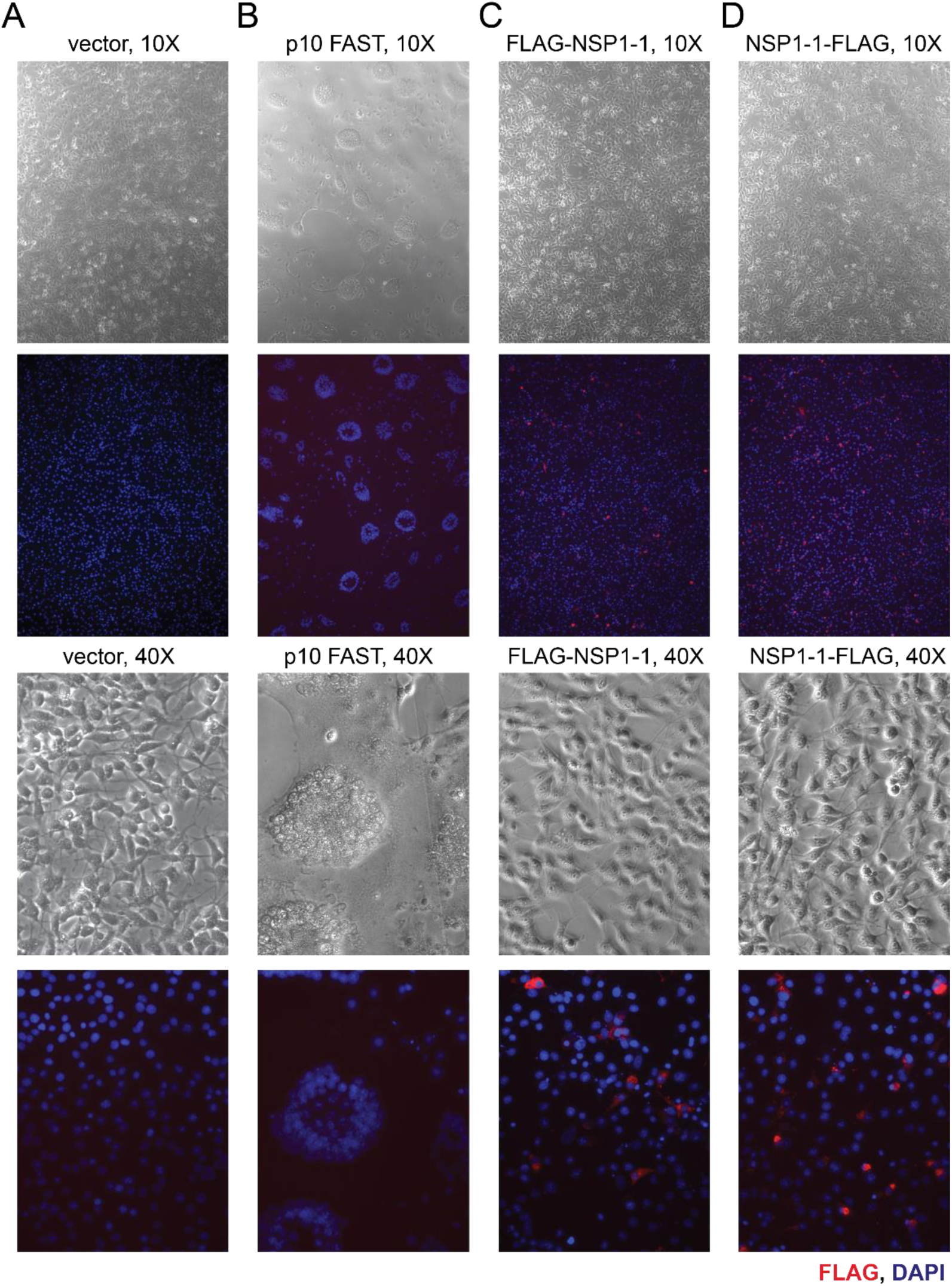
RVB NSP1-1 fails to mediate fusion and exhibits perinuclear localization in BHK cells. BHK-T7 cells were transfected with vector alone (A) or plasmids encoding NBV p10 FAST (B) or RVB Bang117 NSP1-1 with an N-terminal (C) or C-terminal (D) FLAG tag. Cells were fixed and stained with antibodies to detect FLAG (red) or nuclei (blue) and imaged using both differential interference contrast and immunofluorescence microscopy. Representative images are shown. Plasmids used for transfection and objective lens magnification are indicated.

### RVB NSP1-1 mediates enhanced RVA replication in human cells

Since NSP1-1 did not mediate cell-cell fusion in BHK-T7 cells, we sought to test RVB NSP1-1 function in the context of viral infection in Caco-2 and 293T cells, which can support RVA replication and RVB NSP1-1-mediated fusion. Although the precise mechanism is unknown, NBV p10 has been shown to enhance replication of simian RVA strain SA11 in cells adsorbed at low multiplicity of infection (MOI) during a 16 h time course, which is less than the time required to complete a round of replication and initiate infection of a new subset of cells (32). Trypsin, which cleaves viral attachment protein VP4, activates RVA for optimal infectivity (1). To determine whether NBV p10 or RBV NSP1-1 affected viral replication in Caco-2 or 293T cells, we transfected these cells with increasing concentrations of pCAGGS alone or pCAGGS encoding NBV p10 or RVB NSP1-1, then infected them with trypsin-activated RVA strain SA11. At 16 h post infection, we lysed the cells and quantified viral titers. We found that SA11 titer was enhanced in Caco-2 cells transfected with 2 µg of NBV P10 or 1 or 2 µg of NSP1-1, in comparison to vector-transfected cells (Fig. 6A). A very modest but statistically significant enhancement of SA11 titer also was detected in 293T cells transfected with 2 or 10 ng of pCAGGS expressing RVB NSP1-1, in comparison to vector-transfected cells (Fig. 6B). These findings suggest a potential role for RVB NSP1-1 in enhancing rotavirus replication in human cells during a single infectious cycle.

**Figure 6.**
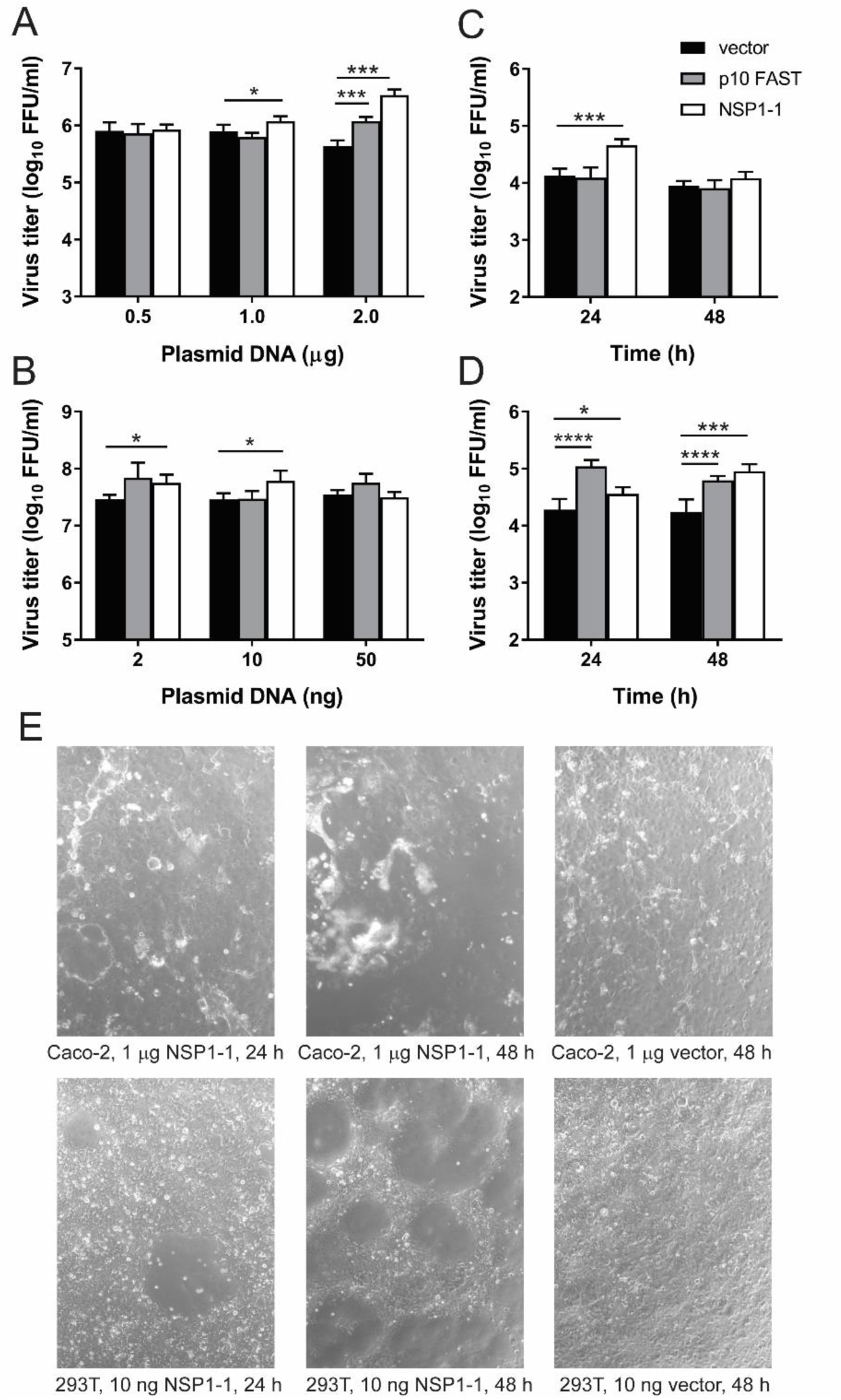
RVB Bang117 FAST mediates enhanced rotavirus replication in human cells. (A-B) Short-term infection. Caco-2 cells (A) or 293T cells (B) were transfected with the indicated amount of plasmid DNA. At 4 h post transfection, cells were adsorbed with trypsin-activated SA11 rotavirus, washed, incubated in serum-free medium containing trypsin at 37°C for 16 h, and lysed. Virus titers in cell lysates were quantified by FFA. (C-D) Rotavirus spread in the presence of FBS. Caco-2 cells were transfected with 1 µg of plasmid DNA (C), or 293T cells were transfected with 10 ng of plasmid DNA (D). At 4 h post transfection, cells were adsorbed with trypsin-activated SA11 rotavirus, washed, incubated in MEM containing 20% (Caco-2) or 10% (293T) FBS for 0, 24, or 48 h, and lysed. Virus titers in cell lysates were quantified by FFA. *n* = 3. *, *p* <0.05; ***, *p* <0.001; ****, *p* <0.0001 in comparison to vector alone by unpaired *t* test. (E) Cytopathic effects in transfected cells. Caco-2 or 293T cells were transfected with the indicated concentrations of plasmids and imaged using brightfield microscopy at 24 or 48 h post transfection to reveal gross morphological changes in the monolayer.

Rotavirus spreads poorly in cultured cells in the presence of fetal bovine serum (FBS), likely due to inhibited cleavage of the viral attachment protein. To determine whether RVB NSP1-1 could facilitate rotavirus spread in the presence of FBS, we transfected Caco-2 or 293T cells with pCAGGS alone or pCAGGS encoding NBV p10 or RVB NSP1-1, then infected them with RVA strain SA11 and incubated them in standard culture medium containing 20% FBS (Caco-2) or 10% FBS (293T). At 24 h and 48 h post transfection, we lysed the cells and quantified viral titer. In Caco-2 cells, we found a modest increase in SA11 titer (<10-fold) at 24 h post infection in RVB NSP1-1 transfected cells in comparison to vector-transfected cells (Fig. 6C). No significant difference in titer was detected at 48 h post infection. However, by this time, RVB NSP1-1-transfected Caco-2 monolayers displayed evidence of significant cytopathic effects, including cell rounding and lifting, which may indicate poor cell health (Fig. 6E). In 293T cells, we found that SA11 titers were significantly enhanced for both NBV p10- and RVB NSP1-1-transfected cells, in comparison to vector-transfected cells at 24 and 48 h post infection (Fig. 6D). Transfection of 293T cells with similar amounts of pCAGGS expressing RVB NSP1-1 results in modest (24 h) to significant (48 h) visible syncytium formation within the monolayer, without complete monolayer disruption and cell lifting (Fig. 6E). Together, these findings suggest that RVB NSP1-1 can enhance rotavirus replication during multi-cycle infection, perhaps by enabling cell-cell spread.

## DISCUSSION

Based on our findings, we propose that RVB encode functional FAST proteins that contain a myristoyl moiety on the N terminus, an extracellular N-terminal loop, a central transmembrane helix, and a relatively short cytoplasmic tail containing a region of approximately four basic residues (Fig. 3C). Short stretches of hydrophobic residues also are predicted in either the N- or C-terminal regions of the protein. The morphological changes induced in cultured cells following NSP1-1 expression, which include the appearance of smooth patches lacking distinct cell membranes and resemble those induced by NBV p10, suggest that RVB NSP1-1 can mediate syncytia formation in human 293T cells (Fig. 1B). The detection of FLAG-positive multinucleated clusters in NSP1-1-FLAG transfected 293T and Caco-2 cells suggests that the cells expressing NSP1-1 are fusing to one another, and addition of a peptide to the C terminus does not disrupt fusion activity (Figs. 2 and 4). The reduced number of clusters in FLAG-NSP1-1-transfected 293T and Caco-2 cells suggests that the N terminus plays an important role in fusion and is consistent with disruption of the N-terminal myristoyl moiety (Figs. 2-4). While the proposed sequence and structural features of RVB NSP1-1 remain to be biochemically and structurally validated, sequence alignment and analysis support our model of RVB NSP1-1 organization (Fig. 3). For example, the presence of a predicted myristoylation motif at the N terminus of every RVB, RVG, and RVI sequence in GenBank (Figs. 3B and S1) provides support for the presence of this fatty acid modification.

If our model is correct, RVB NSP1-1 proteins are the shortest and simplest proteins shown to mediate functional cell-cell fusion. Conservation of a fatty acid modification at the N terminus and the polybasic motif in the cytoplasmic tail in all NSP1-1 proteins, as well as clusters of hydrophobic residues in either the N- or C-terminal domain, suggests these motifs may be critical for protein function (Figs. 3B-C and S1). The myristoylated N terminus and hydrophobic patch residues of FAST proteins induce lipid mixing between liposomal membranes (37, 40). While insertion of N-terminal fatty acid and hydrophobic moieties may promote membrane merger, cytoplasmic hydrophobic patches may promote pore formation by partitioning into the curved rim of a newly formed fusion pore (31). Sequence alignments suggest that RVG and RVI also may encode FAST proteins (Fig. 3B), though their functionality remains to be tested. It will be informative to determine whether these proteins can mediate cell-cell fusion in the absence of evident hydrophobic patches and polyproline motifs. If functional, RVI NSP1-1 will represent the smallest known FAST protein and may help define the minimal requirements for cell-cell fusion. Expansion of the repertoire of known viral FAST proteins may enable the establishment of guidelines that permit identification of additional novel viral FAST proteins, despite their highly divergent sequences.

Many viruses exhibit tropism for certain animal species or cell types. Zoonotic transmission, however, is a frequent but often incompletely understood phenomenon. Virus and host factors can serve as barriers to transmission or virulence factors following zoonotic transmission. Based on the observation that it can mediate fusion in human cells but not hamster cells, NSP1-1 may serve as a viral tropism determinant. However, cells from many different animal species and tissue types, as well as NSP1-1 proteins from rotaviruses derived from different animal hosts, will need to be tested before the boundaries of this limitation are revealed. Support for the idea that there are host species preferences comes from the observation that NSP1 gene sequences of RVBs from murine, human, ovine, bovine, and porcine hosts cluster in distinct phylogenetic groups, with low identities between them and the greatest diversity detected among porcine RVBs (26). Why NBV p10 FAST is capable of mediating fusion in BHK-T7 cells, while RVB NSP1-1 is not, currently is unclear. While RVB NSP1-1 is predicted to be myristoylated at the N terminus, p10 FAST proteins form an extracellular cysteine loop and are not myristoylated but are palmitoylated at a membrane-proximal site in the N terminus (34, 41-43). These differences in acylation may affect functionality. Based on apparent mislocalization of NSP1-1, it is possible that signals that mediate trafficking from the endoplasmic reticulum, where FAST proteins are translated, to the plasma membrane, via the secretory pathway (38, 44-47), fail to function appropriately in some non-homologous hosts. Chimeric FAST proteins containing individual domain exchanges between RVB Bang117 NSP1-1 and NBV p10, similar to those engineered by Eileen Clancy for other FAST proteins (30, 31, 48), may provide insight into protein domains responsible for the species-specific fusion activity of RVB Bang117 NSP1-1.

Expression of NSP1-1 during infection has not been shown. A direct attempt to detect expression of a product from the NSP1-1 ORF following *in vitro* transcription and translation from full-length IDIR gene 7 and immunoprecipitation with convalescent rat serum proved unsuccessful (29). In the same publication, the authors predicted efficient NSP1-2 translation and much less efficient NSP1-1 translation based on nucleotide sequences flanking the START codons (29). However, villous epithelial syncytial cell formation has been observed in the ileum and jejunum of RVB IDIR-infected neonatal rats, with syncytial cells reported to contain large numbers of viral particles (49, 50). Additionally, an ovine RVB strain produced RVB-positive syncytia on MA104 monolayers (51). In numerous studies of RVB in pigs and cows, researchers have failed to note detection of syncytia, though they may not have been looking for such events. However, in a rodent model of infection with a fusogenic pteropine orthoreovirus (PRV), authors failed to detect syncytia in infected lung tissue when specifically looking for these cells (33). In the case of RVB infection, it has been suggested that syncytia are rapidly sloughed from the intestinal epithelium and therefore easily missed (51). These observations suggest that NSP1-1 is expressed during RVB infection, but low levels of expression and cytotoxic effects may render this protein, and the syncytia whose formation it mediates, difficult to detect.

Our results with cells transiently transfected with plasmids expressing RVB NSP1-1 and infected with RVA SA11 suggest potential functions for NSP1-1 in rotavirus replication and spread (Fig. 6). Since only a single successful *in vitro* culture system has been published for RVB (52), with no follow-up studies, we used RVA to study NSP1-1 function in the context of viral infection. Our experimental results support a role for RVB NSP1-1 in enhancing rotavirus replication in the presence of trypsin at a time point less than the length of a single infectious cycle (Fig. 6A-B) and in the presence of inhibitory FBS at time points that would permit multiple rounds of replication (Fig. 6C-D). The latter result is consistent with enhancement of replication by permitting the virus to spread from cell-to-cell without having to initiate infection at the plasma membrane, whereas the former result suggests another mechanism of replication enhancement. A major drawback to our experiments using RVA is the inability to ensure that NSP1-1 expression and viral infection occurred in the same cell. With a small percentage of cells infected and only a subset of cells transfected with plasmid DNA, it is likely that only a fraction of infected cells also fused to adjacent cells to mediate virus spread. In our system, we also were unable to modulate NSP1-1 expression levels, and our codon-optimized expression construct may have yielded significantly higher levels of protein expression than are likely to occur during natural infection, fusing cells too rapidly and resulting in cell death (Fig. 6E). Ultimately, these preliminary observations will need to be validated in a more biologically relevant system. Our results are consistent with those from other published studies showing that FAST proteins can enhance replication of dsRNA viruses on sub-single-cycle and multi-cycle time scales (32, 33). In one study, the authors detected enhancement of viral RNA synthesis in the presence of FAST proteins as early as five hours post infection, and they hypothesized that cell-to-cell fusion provides access to additional substrates for viral transcription, such as nucleotide triphosphates and S-adenosyl methionine (33). Since replication enhancement conferred by FAST proteins was detected even at high MOI, the authors suggested that enhancement is not mediated by cell-to-cell spread. Mechanisms by which cell fusion could enhance viral replication at high MOI remain unclear, as fusion would not be anticipated to provide access to new sources of material for building progeny virions under these conditions.

What is the biological function of RVB NSP1-1 during infection? Syncytia formed between epithelial cells may increase rates of cell-to-cell spread and enhancing viral replication and shedding within the infected host. This hypothesis is supported by the previous detection of syncytial cells, which were reported to contain the majority of virus particles, at the tips of jejunal and ileal villi during RVB infection of neonatal rats (49, 50). Close cellular apposition, mediated by adherens junctions, facilitates FAST protein-mediated fusion (53). Thus enterocytes, which form close contacts via tight junctions, are ideal candidate cells for FAST protein-mediated syncytium formation. While the hypothesis remains to be tested, our findings (Fig. 6C-D) suggest the possibility that cell-to-cell fusion induced by NSP1-1 aids in the spread of RVB strains following introduction into the gastrointestinal tract. Such a mechanism could potentially enable immune evasion within the host by permitting viral spread in the presence of neutralizing antibodies. A report of dairy cows involved in a 2002 RVB diarrhea outbreak shedding a highly similar strain of RVB during a 2005 outbreak suggests that animals are not completely protected after the initial infection (54), though RVA reinfection with reduced disease severity also occurs (1, 55). Regardless of mechanism, there is now published evidence supporting a biological role for a FAST protein *in vivo* (33). In a rodent model, two PRV viruses that are isogenic except in the capacity to express p10 FAST exhibited significant differences in pathogenesis, with animals infected with PRV containing an intact FAST protein exhibiting reduced body weight and survival and enhanced viral titer and lung pathology, in comparison to animals infected with PRV lacking p10 FAST expression. While we currently lack a system in which to directly test its function, these observations suggest a potential role for RVB NSP1-1 and other FAST proteins in viral replication and pathogenesis *in vivo*.

Although it is reasonable to anticipate that NSP1-1 permits evasion of adaptive immune responses by promoting direct rotavirus cell-to-cell spread within the host, it is unclear how RVB and other viruses in its clade (RVG, RVH, and RVI) evade innate immune signaling in the absence of an RVA NSP1 homolog. RVA NSP1 has been shown to promote degradation of innate signaling molecules, including interferon regulatory factors and β-TrCP (3, 56). In some cases, RVA NSP1 function is host species-specific. Perhaps cell-cell fusion permits rotavirus to evade some innate immune mechanisms, or perhaps NSP1-2, whose function remains unknown, obviates the need for an RVA-like NSP1 protein. The evolutionary mechanisms through which FAST proteins became incorporated into rotavirus genomes, or *Reoviridae* genomes in general, and consequences of the lack of an NSP1-1 ORF for RVH viruses also are unclear (57). Future studies using new animal models and technologies, such as reverse genetics and human intestinal organoid culture (32, 58), may permit insights into differences in host interactions among the rotavirus species.

## MATERIALS AND METHODS

### Cells, viruses, and antibodies

Human embryonic kidney 293T cells were grown in Dulbecco’s modified Eagle’s minimal essential medium (Corning) supplemented to contain 10% fetal bovine serum (FBS) (Gibco) and 2 mM L-glutamine. Human colonic epithelial Caco-2 cells were grown in Eagle’s minimum essential medium (MEM) with Earle’s salts and _L_-glutamine (Corning) supplemented to contain 20% FBS, 1X MEM non-essential amino acids (Sigma), 10 mM HEPES (Corning), and 1 mM sodium pyruvate (Gibco). Monkey kidney epithelial MA104 cells were grown in MEM with Earle’s salts and _L_-glutamine (Corning) supplemented to contain 5% FBS. Baby hamster kidney cells expressing T7 RNA polymerase under control of a cytomegalovirus promoter (BHK-T7) (59) were grown in Dulbecco’s modified Eagle’s minimal essential medium (Corning) supplemented to contain 5% fetal bovine serum, 2 mM _L_-glutamine, and 10% tryptose phosphate broth (Gibco). These cells were propagated in the presence of 1 mg/ml G418 (Invitrogen) during alternate passages.

Simian rotavirus laboratory strain SA11 was propagated in MA104 cells, and viral titers were determined by FFA using MA104 cells (60).

Monoclonal mouse anti-FLAG antibody (Sigma), sheep polyclonal rotavirus antiserum (Invitrogen), Alexa Fluor 546-conjugated anti-mouse IgG (Invitrogen), and Alexa Fluor 488-conjugated anti-sheep IgG (Invitrogen) are commercially available.

### Plasmids

NBV p10 in pCAGGS has been described previously (32). pLIC8 was constructed by engineering a ligation-independent cloning site in mammalian expression plasmid pGL4.74. pLIC6 was constructed by engineering a ligation-independent cloning site into mammalian expression plasmid pCAGGS. RVB Bang117 was sequenced from a specimen obtained in 2002 from a 32 year-old male with severe diarrhea in Bangladesh (61). A codon-optimized version of RVB Bang117 NSP1-1 was synthesized (Genscript) and cloned into pLIC8 and pLIC6 using ligation-independent cloning following PCR amplification with appropriate primers and T4 DNA polymerase treatment. Tagged versions of NSP1-1 were engineered using ‘round the horn PCR. Briefly, a pair of primers, each encoding half of the FLAG peptide (DYKDDDDK), was used to amplify NSP1-1 in pLIC8. Then, the PCR fragment was purified and ligated to form a complete tag inserted at the N or C terminus of the ORF. After verifying the sequences of plasmid clones, tagged versions of NSP1-1 were transferred into pLIC6 using ligation-independent cloning, and nucleotide sequences were verified by Sanger sequencing.

### Cell Transfection and Imaging

For differential interference contrast imaging, 293T cells (∼5 × 10^5^ per well) in 12-well plates were transfected with 0.2 µg of plasmid DNA per well using LyoVec transfection reagent (InvivoGen), according to manufacturer instructions, incubated at 37°C for 24 h, and imaged using a Zeiss Axiovert 200 inverted microscope. For confocal imaging, glass coverslips were sterilized, coated with poly-L-lysine (Sigma), rinsed, and dried. 293T cells (1.25 × 10^5^ per well) were seeded onto coverslips in 24-well plates and incubated at 37°C one day prior to transfection with 0.1 µg of plasmid DNA per well using LyoVec. At 24 h post transfection, cells were fixed with cold methanol and blocked with PBS containing 1% FBS. FLAG peptides were detected with mouse anti-FLAG diluted 1:500, and Alexa Fluor 546-conjugated anti-mouse IgG, diluted 1:1000, and nuclei were detected using 300 nM 4′,6-diamidino-2-phenylindole (DAPI, Invitrogen), with washes in PBS containing 0.5% Triton X-100 (Fisher Scientific). Stained coverslips were mounted on glass slides using ProLong Gold antifade mountant (Invitrogen) and dried prior to imaging with an Olympus FV-1000 Inverted confocal microscope.

BHK-T7 cells (∼2 × 10^5^ per well) in 24-well plates were transfected with 0.5 µg of plasmid DNA per well using TransIT-LT1 transfection reagent (Mirus Bio) in OptiMEM (Gibco), according to manufacturer instructions, and incubated at 37°C for 24 h prior to fixing and staining. Cells were fixed with cold methanol and blocked with PBS containing 1% FBS. Staining to detect FLAG and nuclei was performed as described above. Stained cells were imaged using a Zeiss Axiovert 200 inverted microscope equipped with an HBO 100 mercury arc lamp.

Caco-2 cells (∼1 × 10^5^ per well) in 24-well plates were transfected with 1 µg per well of plasmid DNA using TransIT-LT1 transfection reagent in OptiMEM, according to manufacturer instructions, and incubated at 37°C for 24 h prior to fixing and staining. Cells were fixed with 10% neutral buffered formalin and blocked with PBS containing 0.5% Triton X-100 and 5% FBS. Staining to detect FLAG and nuclei was performed as described above. Stained cells were imaged using a Zeiss Axiovert 200 inverted microscope equipped with an HBO 100 mercury arc lamp.

### Quantitation of single cells, clusters, and cluster diameter

Caco-2 cells in 24-well plates were transfected with 1 µg per well of plasmids encoding RVB Bang117 NSP1-1-FLAG or RVB Bang117 FLAG-NSP1-1 and stained to detect FLAG and nuclei as described for imaging studies. Clusters were defined as groupings of at least three FLAG-positive cells in contact with one another. To quantify single cells and clusters, entire wells of transfected cells were visually analyzed using a Zeiss Axiovert 200 inverted microscope. The person analyzing the wells was not the person who performed the transfections and in most cases was blinded to the identity of the samples. Four independent experiments each containing three technical replicates were analyzed. To quantify cluster diameter, entire wells of transfected cells from the four independent experiments just described were imaged using an ImageXpress Micro XL automated microscope imager (Molecular Devices). Diameters of 20 cell clusters per plasmid construct per experiment were quantified using MetaXpress image analysis software v6.5 (Molecular Devices). To compare numbers of clusters, single cells, and cluster diameters, statistical analyses were performed using unpaired t-tests with GraphPad Prism 7 (GraphPad).

### Transfection-Infection Experiments in Caco-2 cells

For short-term rotavirus transfection-infection experiments, Caco-2 cells (∼1 × 10^5^ per well) in 24-well plates were transfected with 0.5, 1, or 2 µg of plasmid DNA per well using TransIT-LT1 transfection reagent in OptiMEM and incubated at 37°C for 3 h. Medium was removed from the transfected cells and replaced with serum-free MEM for 1 h prior to virus adsorption. SA11 rotavirus was activated by incubation with 1 µg/ml trypsin at 37°C for 1 h. Medium was removed from the cells, and they were adsorbed with activated SA11 rotavirus diluted in 0.1 ml of serum-free MEM per well to a MOI of 0.1 PFU/cell at 37°C for 1 h. After adsorption, cells were washed then incubated in serum-free MEM containing 0.5 µg/ml of trypsin at 37°C for 16 h. Cell lysates were harvested after three rounds of freezing and thawing, and virus in the resultant lysates was quantified by FFA.

For longer-term rotavirus transfection-infection experiments, Caco-2 cells (∼1 × 10^5^ per well) in 24-well plates were transfected with 1 µg of plasmid DNA per well using TransIT-LT1 transfection reagent in OptiMEM and incubated at 37°C for 3 h. Medium was removed from the transfected cells and replaced with serum-free MEM for 1 h prior to virus adsorption. SA11 rotavirus was activated by incubation with 1 µg/ml trypsin at 37°C for 1 h. Medium was removed from the cells, and they were adsorbed with activated SA11 rotavirus diluted in 0.1 ml of serum-free MEM per well to a MOI of 1 PFU/cell at 37°C for 1 h. After adsorption, cells were washed then incubated in MEM containing 20% FBS at 37°C for 24 or 48 h. Cell lysates were harvested after three rounds of freezing and thawing, and virus in the resultant lysates was quantified by FFA.

### Transfection-Infection Experiments in 293T cells

For short-term rotavirus transfection-infection experiments, 293T cells (∼2.5 × 10^5^ per well) in 24-well plates were transfected with 50, 10, or 2 ng of plasmid DNA per well using LyoVec and incubated at 37°C for 3 h. Medium was removed from the transfected cells and replaced with serum-free DMEM for 1 h prior to virus adsorption. SA11 rotavirus was activated by incubation with 1 µg/ml trypsin at 37°C for 1 h. Medium was removed from the cells, and they were adsorbed with activated SA11 rotavirus diluted in 0.1 ml of serum-free DMEM per well to a MOI of 1 PFU/cell at 37°C for 1 h. After adsorption, cells were washed then incubated in serum-free DMEM containing 0.5 µg/ml trypsin at 37°C for 16 h. Cell lysates were harvested after three rounds of freezing and thawing, and virus in the resultant lysates was quantified by FFA.

For longer-term rotavirus transfection-infection experiments, 293T cells (∼2.5 × 10^5^ per well) in 24-well plates were transfected with 6 ng of plasmid DNA per well using LyoVec and incubated at 37°C for 3 h. Medium was removed from the transfected cells and replaced with serum-free DMEM for 1 h prior to virus adsorption. SA11 rotavirus was activated by incubation with 1 µg/ml trypsin at 37°C for 1 h. Medium was removed from the cells, and they were adsorbed with activated SA11 rotavirus diluted in 0.1 ml of serum-free DMEM per well to a MOI of 0.1 PFU/cell at 37°C for 1 h. After adsorption, cells were washed then incubated in DMEM containing 10% FBS at 37°C for 24 or 48 h. Cell lysates were harvested after three rounds of freezing and thawing, and virus in the resultant lysates was quantified by FFA.

### Fluorescent focus assay

MA104 cells (4 × 10^5^ per well) were seeded in black-wall 96-well plates and incubated overnight until near confluency. Infected cell lysates were activated with 1 µg/ml trypsin for at 37°C for 1 h then serially diluted in serum-free MEM. Medium was removed from MA104 cells, they were washed twice in serum-free MEM and adsorbed with serial virus dilutions at 37°C for 1 h. Inocula were removed, cells were washed with serum-free MEM then incubated in fresh medium at 37°C for 14-18 h. Cells were fixed with cold methanol, and rotavirus proteins were detected by incubation with sheep polyclonal rotavirus antiserum at a 1:500 dilution in PBS containing 0.5% Triton X-100 at 37°C, followed by incubation with Alexa Fluor 488-conjugated anti-sheep IgG diluted 1:1000 and 300 nM DAPI. Images were captured for four fields of view per well using an ImageXpress Micro XL automated microscope imager (Molecular Devices). Total and percent infected cells were quantified using MetaXpress high-content image acquisition and analysis software (Molecular Devices). Fluorescent foci from four fields of view in duplicate wells for each sample were quantified. Statistical analyses were performed using GraphPad Prism 7 (GraphPad). Virus titers in cells transfected with plasmids expressing NBV p10 or RBV NSP1-1 were compared to those in vector-transfected cells using an unpaired t-test.

### Amino acid alignments and phylogenetic analysis

Sequences of NSP1-1 were obtained from GenBank. Accession numbers for FAST sequences analyzed for the ML tree shown in Fig. 3A and alignment in Fig. 3B are ACN38055, ADZ31982, ABV01045, AAM92750, AAM92738, AAF45151, ABY78878, AAF45157.1, ABM67655, ACU68609, AAP03134, AHL26969, and AAL01373. Accession numbers for rotavirus segment 5 source sequences for NSP1-1 proteins analyzed for the ML tree shown in Fig. 3A are KY689691, NC_021546, KX362373, KC876005, NC_021583, NC_026820, and KY026790. In each case, the smaller ORF sequence was analyzed. Accession numbers for rotavirus segment 5 source sequences for NSP1-1 proteins in Figs. 3B and S1 are indicated.

For phylogenetic analysis, amino acid sequences were aligned using the MUSCLE algorithm in MEGA 7.0 (62). The Le Gascuel 2008 model (63) was selected as the best-fit model by Modeltest and used in maximum likelihood (ML) phylogeny construction with 1000 bootstrap replicates (in MEGA 7.0). Initial tree(s) for the heuristic search were obtained automatically by applying Neighbor-Join and BioNJ algorithms to a matrix of pairwise distances estimated using a JTT model, and then selecting the topology with superior log likelihood value. A discrete Gamma distribution was used to model evolutionary rate differences among sites (5 categories (+G, parameter = 8.9228)). The tree with the highest log likelihood is shown in Fig. 3A. The ML tree was visualized using FigTree v1.4.2 (http://tree.bio.ed.ac.uk/software/figtree/).

For Figs. 3B and S1, amino acid alignments were constructed with MAFFT v7.2 (64) using the E-INS-I strategy. N-myristylation motifs were defined using ScanProsite (65). Transmembrane helices were predicted with the TMHMM Server v2.0 (www.cbs.dtu.dk/services/TMHMM/). Hydrophobic, polybasic, and polyproline regions for FAST proteins were identified previously (31). Hydrophobic patches for NSP1-1 proteins were identified using ProtScale, with a window size of nine amino acids (66). Polybasic regions for NSP1-1 proteins were identified visually.

## ACKNOWLEDGEMENTS

We thank Dr. Takeshi Kobayashi for NBV p10 in pCAGGS and Dr. Marco Morelli for pLIC6 and pLIC8 plasmids.

This research was supported in part by CTSA award No. UL1 TR002243 from the National Center for Advancing Translational Sciences. Its contents are solely the responsibility of the authors and do not necessarily represent official views of the National Center for Advancing Translational Sciences or the National Institutes of Health.

---

**Figure S1.**
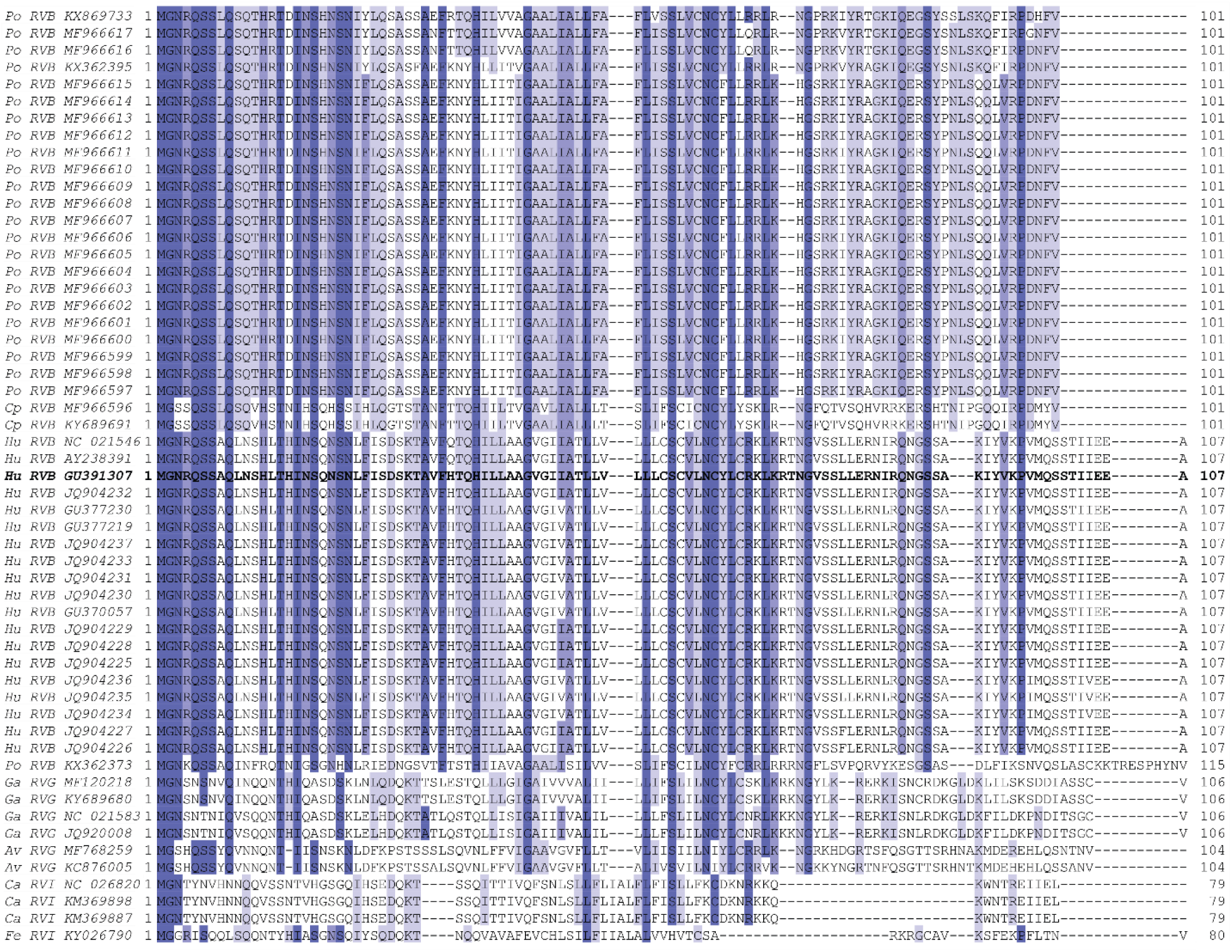
Alignment of complete RVB, RVG, and RVI NSP1-1 sequences, colored based on amino acid identity, with darker purple indicating higher identity at a given position. RVB Bang117 NSP1-1 is shown in bold text. For each sequence, host origin, rotavirus species, and GenBank accession number are indicated. Av, avian; Ca, canine; Cp, caprine; Fe, feline; Ga, gallinaceous; Hu, human; Po, porcine.

